# Patient-derived tumor explant models of tumor immune microenvironment reveal distinct and reproducible immunotherapy responses

**DOI:** 10.1101/2024.05.11.593502

**Authors:** Rita Turpin, Karita Peltonen, Jenna H. Rannikko, Ruixian Liu, Anita N. Kumari, Daniel Nicorici, Moon Hee Lee, Minna Mutka, Panu E. Kovanen, Laura Niinikoski, Tuomo Meretoja, Johanna Mattson, Petrus Järvinen, Kanerva Lahdensuo, Riikka Järvinen, Sara Tornberg, Tuomas Mirtti, Pia Boström, Ilkka Koskivuo, Anil Thotakura, Jeroen Pouwels, Maija Hollmén, Satu Mustjoki, Juha Klefström

## Abstract

Tumor-resident immune cells play a crucial role in eliciting anti-tumor immunity and immunomodulatory drug responses, yet these functions have been difficult to study without tractable models of tumor immune microenvironment (TIME). Patient-derived *ex vivo* models contain authentic resident immune cells and therefore, could provide new mechanistic insights into how TIME responds to tumor or immune cell-directed therapies. Here, we assessed the reproducibility and robustness of immunomodulatory drug responses across two different *ex vivo* models of breast cancer TIME and one of renal cell carcinoma. These independently developed TIME models were treated with a panel of clinically relevant immunomodulators, revealing remarkably similar changes in gene expression and cytokine profiles among the three models in response to T cell activation and STING-agonism while still preserving individual patient-specific response patterns. Moreover, we found two common core signatures of adaptive or innate immune responses present across all three models and both types of cancer, potentially serving as a benchmark for drug-induced immune activation in *ex vivo* models of TIME. The robust reproducibility of immunomodulatory drug responses observed across diverse *ex vivo* models of TIME underscores the significance of human patient-derived models in elucidating the complexities of antitumor immunity and therapeutic interventions.

## INTRODUCTION

The field of immuno-oncology has revolutionized cancer care. Nevertheless, the effectiveness of immunotherapies on solid tumors has been notably modest, a phenomenon largely attributed to the complex interplay between tumor cells and immune-modulating factors within the tumor immune microenvironment (TIME)^1^. A comprehensive examination of the tumor resident immune landscape, both pre- and post-immunotherapeutic interventions, has yielded insights into underlying factors contributing to the unresponsiveness of certain cancer types to immunotherapies^2^. This increasing understanding of the TIME holds immense promise for advancing the frontiers of solid tumor immunity.

Several patient-derived *ex vivo* models have been developed in pursuit of uncovering resident immune cell interactions within the TIME^2–5^. However, uncertainty remains regarding the optimization of *ex vivo* models for immuno-oncology research. On one hand, it is crucial to mimic the natural tumor tissue characteristics to understand interactions that occur under physiological conditions and tumor architecture^6^. However, creating more physiological models embedded in stiff materials and microfluidics devices, for example, may be prohibitively costly, or limit downstream applications. On the other hand, disregarding physiological conditions may induce artificial results and unwanted effects like immune cell efflux in the absence of added extracellular matrix^2^.

To investigate how various culture techniques influence the response of resident immune cells to immunomodulation, distinct *ex vivo* patient-derived explant culture (PDEC) models of breast cancer (BC) and renal cell carcinoma (RCC) were developed independently in three different research groups (**Fig 1A**). These models of breast and renal cell cancer-associated TIME represented increasing layers of processing defined as the manipulation of original tumor architecture and the addition of chemical cues. We used gene expression profiling and cytokine analyses to dissect the response of immunotherapies targeting adaptive and innate immune cells. Our study demonstrates consistent patterns of immune activation across diverse *ex vivo* models of TIME, revealing shared core signatures of adaptive and innate responses to immunomodulatory drugs, irrespective of the specific model or cancer type.

**Figure 1.**
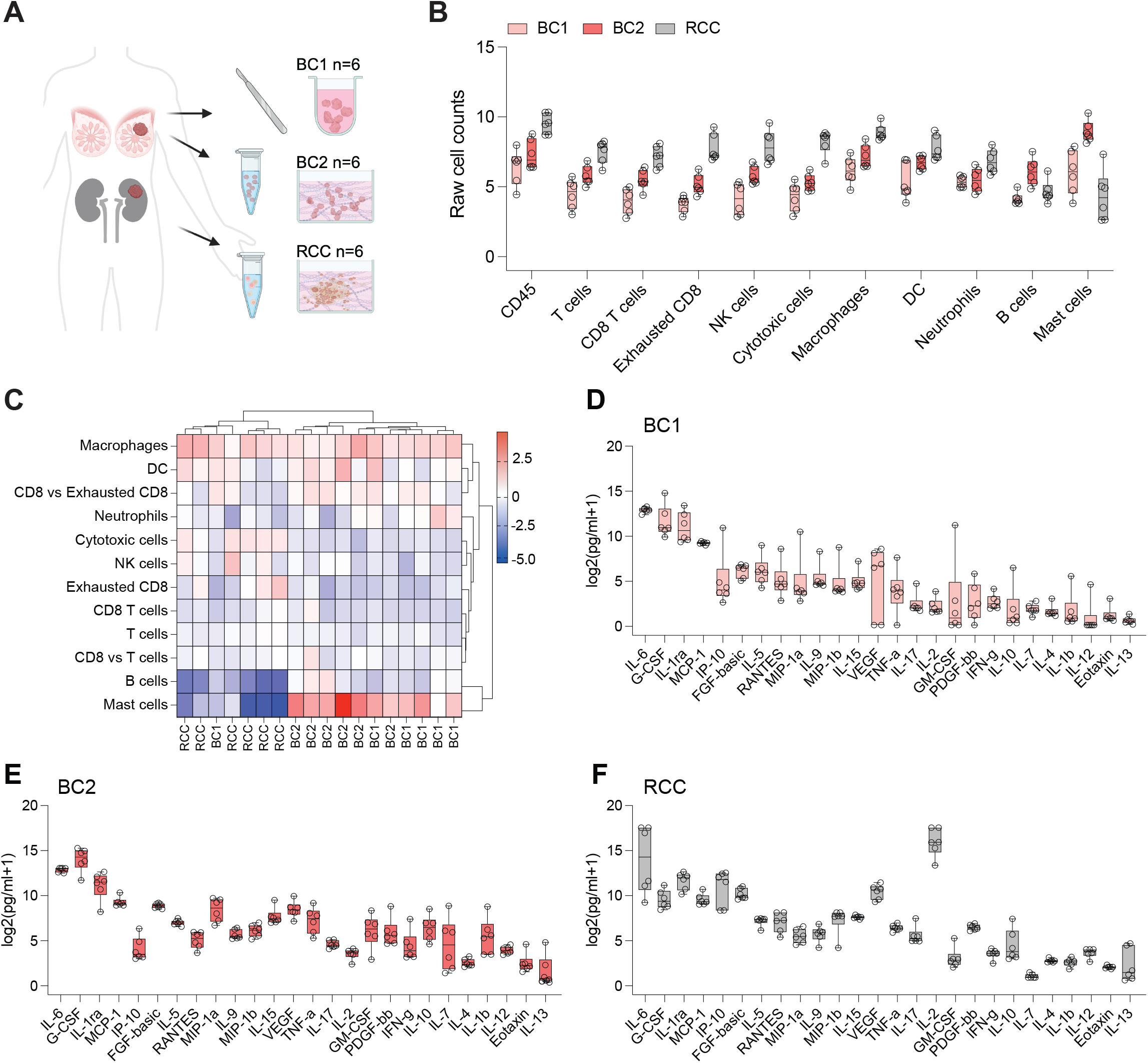
*Ex vivo* tumor landscape of breast cancer and renal cell carcinoma. **A**, Schematic representation of experimental workflow, n=6 for each model, ‘BC1’, ‘BC2’ and ‘RCC’ **B**, Estimated cell type scores (min to max with median) based on gene expression profiling ‘BC1’ = light pink, ‘BC2’ = dark pink and ‘RCC’ = grey **C**, Estimated cell type score per patient normalized to tumor immune infiltration clustered by Ward’s minimum variance method. Scale is z-score. **D-F**, Box and whisker (min to max with median) plot of baseline cytokine secretion of each patient presented as log_2_ of the raw pg/ml (+1 as a small constant value).

## MATERIALS AND METHODS

### Isolation of biological material and patient-derived models

#### PDEC “BC1” minimal manipulation; intact tissue architecture

Tumor tissue samples were collected in RPMI medium (Gibco) from treatment-naïve patients with breast cancer undergoing mastectomy surgery at the Turku University Hospital from Jan 2021 to Aug 2021. Written informed consent was obtained from each participant and the study was conducted under the approval of The Ethics Committee of the Hospital District of Southwest Finland (decision number: ETMK 132/2016) and in accordance with the ethical principles of the declaration of Helsinki. Fresh tumor tissues were cut into small pieces with a diameter of <2 mm and frozen in RPMI (Sigma, cat. R5886) + 10% FCS + 1% GlutaMAX + penicillin-streptomycin (P/S, 12.8 U/mL, Gibco, cat.15140-122) supplemented with 10% DMSO at -150°C. The frozen tissues were thawed in a 37°C water bath and pelleted by centrifugation at 300*g* for 10 min at 10°C. The pieces were further cut (Ø <1.5mm) and transferred on a 96-well low-attachment plate (Corning, cat.7007) for *ex vivo* treatment. The treatments were performed in quadruplicates in 200 μL of RPMI + 10% FCS + 1% GlutaMAX + P/S and three tissue pieces per well.

#### PDEC “BC2” moderate manipulation; partial dissociation and embedding

Fresh tissue was obtained from the treatment-naïve elective breast cancer surgeries performed at the Helsinki University Central Hospital from 2020-2023 (Ethical permit: 243/13/03/02/2013/ TMK02 157 and HUS/2697/2019 approved by the Helsinki University Hospital Ethical Committee). Patients participated in the study by signing an informed consent form.

Explants were made by incubating fresh primary tumor tissue overnight in collagenase A (3 mg/mL; Sigma) in MammoCult media (StemCell technologies) supplemented with MammoCult proliferation supplement (StemCell technologies) with gentle shaking (130 rpm) at 37°C. The resulting explants were collected via centrifugation at 353*g* for 5 min and washed once with PBS. Isolated explants were resuspended in Cultrex Reduced Growth Factor Basement Membrane Extract, Type 2 (R&D Systems). 37.5μL of matrix was pipetted in the center of each well of an 8-chamber slide (Thermo Scientific) in supplemented MammoCult media.

#### PDEC “RCC” high manipulation; single cell dissociation, embedding, and media supplementation

Renal cell carcinoma tumor tissue was obtained from treatment-naïve patients undergoing radical or partial nephrectomy at the HUS Helsinki University Hospital. Studies were approved by the Helsinki University Hospital Ethical Committee (Dnro 115/13/03/02/15) and samples were obtained upon written informed consent. Tissues were preserved in MACS^®^ tissue storage solution during transportation to the laboratory. Upon arrival, tumor tissue was dissociated using Miltenyi’s Tumor Dissociation kit (Miltenyi Biotec), and dissociated cells were live-frozen in 10% DMSO-FBS and stored at -150°C until use.

Frozen tumor dissociates were thawed in AIM-V media, resuspended in Cultrex 3D Culture Matrix BME (Trevigen), and applied on 8-well chamber slides (Lab-Tek). Following matrix solidification, AIM-V media supplemented with recombinant FGF (100 ng/mL), EGF (50 ng/mL), IL-2 (60 IU/mL), sodium pyruvate (10 mM), B-27 (1.5%), and r-spondin (50 ng/mL).

### Drug treatments

Before experimentation, all drugs were aliquoted and distributed to ensure that each research group had access to the exact same lots. Each drug was aliquoted to minimize freezing/thawing cycles. The explants were either left untreated or treated with 25 μL/mL Immunocult (Stemcell, cat.10970), 10 μM ADU-S100 (MedChemExpress, cat. HY-12885), 50 μg/mL pembrolizumab (MedChemExpress, cat. HY-P9902A), 10 μg/mL magrolimab (Icosagen, clone 5F9) or 50 μg/mL pembrolizumab and 10 μg/mL magrolimab. After 48h of *ex vivo* treatment at 37°C and 5% CO_2_, both tissues/cells and culture supernatants were collected and separated by centrifugation for subsequent NanoString and cytokine profiling, respectively.

### NanoString gene expression profiling

BC1 tissue pieces from replicate wells were collected in 1 mL of TRIsure (BioLine, cat. BIO-38032) and homogenized in gentleMACS M tubes (Miltenyi Biotec, cat. 130-093-236) using gentleMACS dissociator (Miltenyi Biotec, cat. 130-093-235) with program RNA.01_01. RNA was extracted according to the manufacturer’s protocol for TRIsure. Shortly, the samples were mixed with 200 μL chloroform and centrifuged at 12’000*g* for 15 min at 4°C. The aqueous phase containing the RNA was collected and precipitated with 500 μL cold isopropyl alcohol for 10 min at RT, and samples were centrifuged at 12’000*g* for 10 min at 4°C. Pellets were washed once with 1 mL 75% ethanol, air-dried, and dissolved in nuclease-free water (Ambion, cat. AM9930). For BC2 and RCC, total RNA was isolated using RNeasy (Qiagen) according to manufacturer instructions. BC1, BC2, and RCC model DNAse removal and RNA concentration were performed using a commercial kit (Zymo Research, cat. R1013) according to the manufacturer’s instructions.

RNA amount was measured with Qubit, and RNA quality was measured with the Agilent Tapestation (BC1 RIN 5.0-9.0; BC2 RIN 5.5-8.2; RCC RIN 6.4-9.5) before gene expression analysis on the NanoString nCounter gene expression platform (NanoString Technologies). nCounter Human PanCancer Immune Profiling Panel consisting of 770 genes identifies different immune cell types and their abundance, measures both the adaptive and innate immunity functions and quantifies checkpoint molecule and cancer-testis (CT) antigen expression. For the supplementary analyses, nCounter PanCancer IO 360 Panel was used, as well. Per sample, 50 ng of total RNA in a final volume of 5 μL was mixed with a reporter codeset, hybridization buffer, and capture codeset. Samples were hybridized at 65°C for 20 hours. Hybridized samples were run on the NanoString nCounter SPRINT profiler.

NanoString gene expression profiling results were analyzed using the nSolver advanced analysis software (v 4.0). During the normalization step, the housekeeping genes with average count of less than 100 were removed. Differential expression analysis expression was performed with Patient_id as a confounding factor. P-values were adjusted by the Benjamini & Yekutieli method^7^. Venn diagrams were plotted from differentially expressed genes with an adjusted P-value of less than 0.05. Pathway scores were calculated as the first principal component of the pathway genes’ normalized expression. Pathway scores were adjusted with Patient_id as a confounding factor. Pathway scores were log_2_-transformed after adding a constant, log_2_FoldChanges calculated from untreated, and resulting scores hierarchically clustered and visualized as a heatmap using R package ComplexHeatmap. The associations between estimated cell abundance in the untreated samples and the fold-changes in immune activity pathways (treated/untreated) were evaluated by computing Pearson correlation. Biomarker evaluation was done using the TIDE^8,9^ biomarker evaluation module where a the average expression of a custom geneset is compared to existing ones in clinical datasets for which survival data is recorded. The module generates corresponding gene expression plots and Kaplan-meier plots.

### Cytokine profiling

Cytokine secretion was analyzed from cleared PDEC culture supernatants using Bio-Plex Pro Human Cytokine 27-plex assay kit (Bio-Rad, cat. M500KCAF0Y) and Bio-Plex 200 System (Bio-Rad) according to the manufacturer’s instructions. Results were analyzed using Bio-Plex Manager 6.0 software (Bio-Rad Laboratories). Cytokines with >10% of data points outside the detection range were excluded from the analyses. The remaining values lower than the detection limit were replaced by 0.5 × lowest measured value. Cytokines were visualized using GraphPad Prism (v.9.5.1). To identify changes in cytokine secretion following treatment, log_2_FoldChanges were calculated.

## RESULTS AND DISCUSSION

### Ex vivo TIME landscape of breast cancer and renal cell carcinoma

Human *ex vivo* tumor models have been used to dissect early immunotherapy responses^2–4^ with promising predictive potential for patients receiving immune-checkpoint blockade (ICB) therapy^2^. However, the development of human-derived *ex vivo* models is not standardized, and it is unclear how the disruption of the tissue architecture and *ex vivo* culture conditions influence immunotherapy responses. Therefore, we tested the reproducibility of the same drug treatments activating innate and adaptive immune cells with three *ex vivo* culture models.

We compared three patient-derived *ex vivo* models in increasing levels of manipulation **(Fig 1A)**. The breast cancer model “BC1” represents the least disruptive system as it has very little hands-on processing, and the tumor architecture is mostly preserved. The breast cancer model “BC2^5^” is fragmented through mild enzymatic digestion also resulting in intact tumor fragments with original tumor architecture^6^. BC2 fragments are further embedded into a 3D matrix, which provides structural support for several cell types. The renal cell carcinoma model “RCC” has the most amount of processing featuring enzymatic digestion of tumor material down to single cells, which are embedded in a 3D matrix and supplemented with IL-2 and growth factors.

Based on gene expression profiling, the baseline immune cell composition of the models differed across cancer types, with RCC being richer in all immune cell types except mast cells and B cells in comparison to the BC models **(Fig 1B)**. This is in line with a reported RNA-based pan-cancer analysis of 33 cancer types, which shows that RCC has more immune infiltrate and is considered more inflammatory^10^. Between the two BC models, BC2 had higher raw immune cell type scores **(Fig 1B)**, possibly reflecting immune efflux in the BC1 model in the absence of matrix embedding, or because the ECM itself provides support for the homeostasis of cell types like dendritic cells (DCs)^11^. Some discrepancies may also arise from the fact that BC1 was frozen prior to culturing. The analysis of the tumor-infiltrating immune cells’ relative abundances revealed that BC1, BC2, and RCC samples had comparable macrophage infiltration, while the BC models had a higher relative proportion of B cells and mast cells, and a lower relative proportion of cytotoxic cells **(Fig 1C)**. Baseline cytokine profiling showed that all models were largely secreting chemokines, for instance, G-CSF, MCP-1, IP-10 (CXCL10), RANTES (CCL5) and MIP-1a/b (CCL3/4) **(Fig 1D-F)**. This is in line with the high myeloid and lymphoid immune cell infiltration within the tumor models. As the cell numbers were not calculated prior to culturing, we assessed cytokine trends rather than cytokine amounts across the models. High levels of the pleiotropic cytokine IL-6 were detected in the TIME of each model. IL-6 may be secreted by various immune cell types as well as cancer cells and its overexpression is reported ubiquitously in many cancer types. High levels of other pro-inflammatory cytokines, such as TNF-α, and interleukins with T-cell and NK-cell stimulatory properties (eg. IL-15) were also present in each model. As expected, RCC samples showed high VEGF secretion. Furthermore, we detected higher IL-2 levels in RCC as a result of IL-2 media supplementation. In comparison to BC1, BC2 had an increased trend towards higher baseline secretion of T helper 2 type cytokines (IL-5, IL-10), proinflammatory cytokines (IL-17, IFN-y, IL-12, IL-1b), and growth factors and regulators (GM-CSF, IL-7, IL-2), possibly reflecting distinct immune landscape and composition in BC2 cultures. G-CSF was generally expressed higher in BC than RCC, perhaps due to the higher relative abundance of neutrophils and mast cells in BC.

Importantly, though individual patient cytokine profiles were consistent for each model type, *ex vivo* modeling did not mask patient-specific variation **(S1A-C)**. For example, “BC1” patient H72 with a clinical diagnosis of triple-negative breast cancer (TNBC) **(S1D)** displayed higher baseline secretion of cytokines like GM-CSF and G-CSF **(S1A)** consistent with previously published findings of high G-CSF secretion by TNBC cells compared to cells of other breast cancer types^12^. Therefore, we found that the baseline immune contexture of each model was concordant with literature for these cancer types, supporting further exploration of differences in immune responses.

### Effects of adaptive and innate immune modulation on immune cell abundance and cytokine secretion

To measure immunotherapy responses in our *ex vivo* models, we selected drugs that target both adaptive and innate immune cells with different modes of action (**Fig 2A**). While anti-PD-1(pembrolizumab) is not widely used in breast cancer, except for TNBC in combination with chemotherapy^13,14^, it shows efficacy in RCC^15–18^ and is accepted as a combination treatment with VEGF/VEGFR inhibition. Magrolimab is a first-in-class humanized monoclonal CD47-binding antibody that prevents tumor cells from escaping macrophage phagocytosis. Magrolimab is being tested in BC in combination with standard therapies (NCT04958785, NCT05807126). CD47 is highly expressed in RCC patients and has been suggested as a therapy target in RCC^19^. Moreover, HX009, a PD-1/CD47 bispecific monoclonal antibody is under phase II clinical trial in solid tumors (NCT04886271), further prompting investigation of pembrolizumab and magrolimab alone and in combination in our models. To achieve maximal immune modulation, we also tested T-cell-activating Immunocult (anti-CD3/CD28/CD2), and the cGAS/stimulator of interferon genes (STING) pathway agonist (ADU-S100) to induce robust adaptive and innate immune activation, respectively.

**Figure 2.**
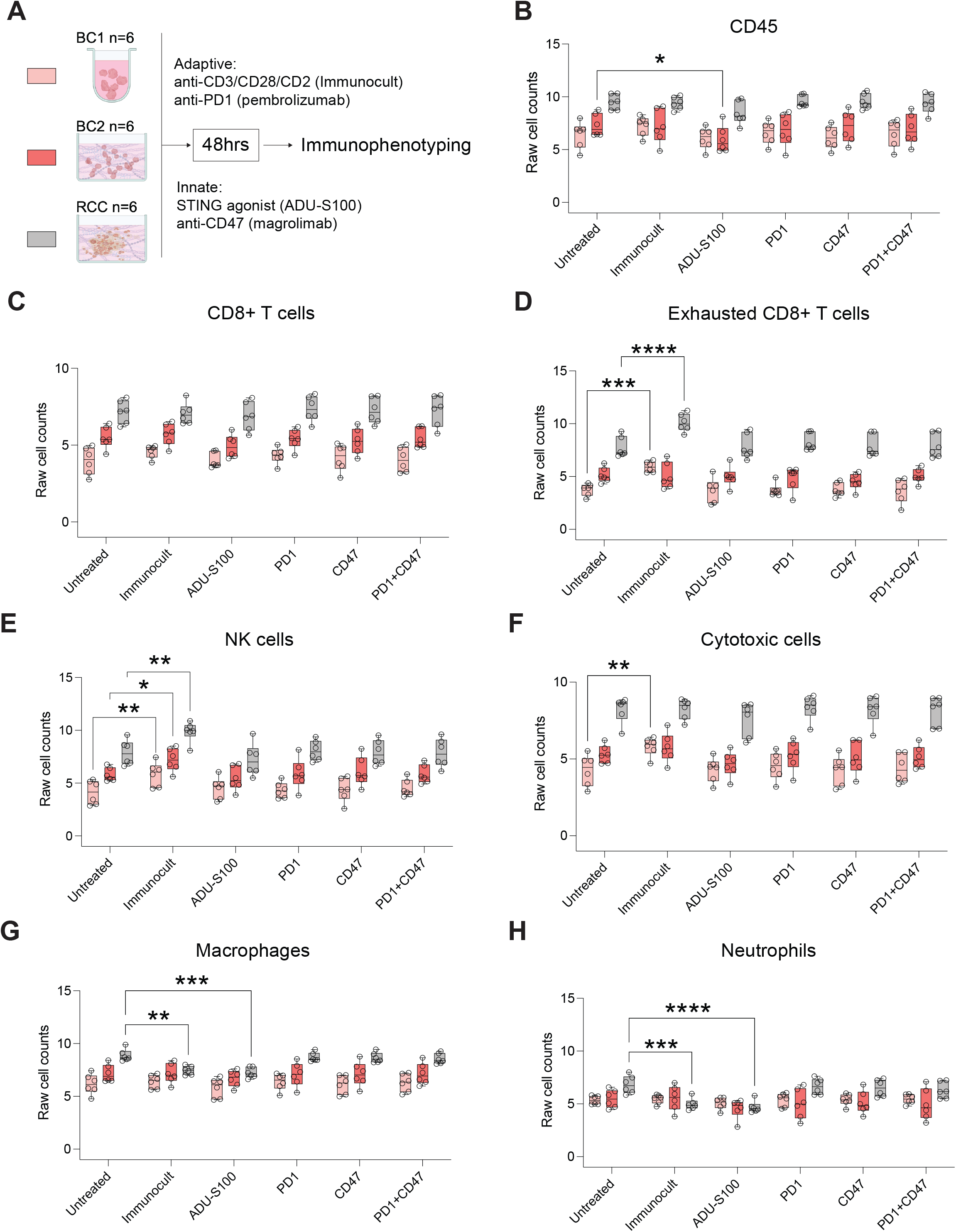
Effect of adaptive and innate immune modulation on cell abundance. **A**, Schematic of treatments selected for the study **B**, Box and whisker (min to max) of total estimated leukocyte abundances after immunomodulation. **C**, No significant changes in CD8+ T cell numbers, but significant changes in **D**, exhausted CD8+ T cells **E**, NK cells **F**, cytotoxic cells **G**, macrophages, and **H**, neutrophils. Statistical significance was tested with a two-way ANOVA with Fishers LSD. All data are presented as min to max box-plot with median. (**** = p <0.0001; *** = p <0.001; ** = p <0.01; * = p <0.05)

Other drug candidates for the study included anti-HER2 (trastuzumab), anti-EGFR (cetuximab), and positive controls lipopolysaccharide (LPS) and a cocktail of IL-2/IL-15. The final drug panel was narrowed down to 6 conditions based on clinical relevance and signal intensity to accommodate samples of two patients on each NanoString cartridge to prevent differences due to batch effects. To prioritize clinical relevance, positive control compounds like LPS and IL-2/IL-15 were mainly included to optimize the readout and timepoints. Selecting drug candidates with adequately robust inflammatory signal induction ensured that there was a strong enough response to compare across all three models. Gene expression profiling (n=1 per treatment) revealed that Immunocult, ADU-S100, LPS and anti-CD47 evoked a stronger response at 48hrs compared to 24hrs **(S2A)**. The remaining treatments showed little effect **(S2E-H)**. Similarly, cytokine profiling of the same samples showed a modest response to anti-PD-1, anti-EGFR, anti-HER2, and IL-2/IL-12 supplementation, although some immune activation was observed, evident by the induction of IP-10 **(S3A)**. We kept anti-PD-1 for its clinical relevance, and discontinued anti-EGFR and trastuzumab due to their weak signal.

Thus, the resulting panel of drugs **(Fig 2A)** was applied for 48hrs and immune cell abundances were studied following treatment. The estimated CD45+ leukocyte abundances were mostly unaffected, aside from a significant decrease in total leukocytes in BC2 following ADU-S100 **(Fig 2B)**. The leukocytes that were depleted in BC2 following ADU-S100 may represent a combination of cell types that were slightly depleted following treatment, but not significantly as individual cell types.

Examples would include T cells and B cells, which were slightly decreased in BC2 **(S4A-B)**. A similar trend was not seen in the other BC model, nor with RCC embedded in the same matrix, suggesting there may be additional cytokines in BC1 and RCC which may support T cells and B cells following STING-agonism. T cells, CD8+ T cells, B cells, DCs, and mast cells did not change in numbers following immunomodulation with any of the tested compounds **(Fig 2C-D, Fig S4A-D)**. However, robust T cell activation achieved with Immunocult did have a significant positive impact on NK cell (BC1, BC2, RCC), cytotoxic CD8+ T cell (BC1), and exhausted CD8+ T cell (BC1, RCC) abundances, and a significant negative effect on macrophages (RCC), and neutrophils (RCC) **(Fig 2D-H)**.

Immunocult induced a robust effect on cytokine secretion, however, the magnitude and the diversity of the response were strikingly different among the models **(Fig 3A-C)**. Cytokines representing immune activation (IP-10 [CXCL10], TNF-α) following T cell stimulation were detected across all three explants. Additionally, BC1 and RCC models embarked significant chemokine (eg. GM-CSF, IP-10 [CXCL10], MIP-1a/b, RANTES, eotaxin) and cytokine (eg. IL-12, IL-10, IL-17, IL-13) secretion. Many samples also showed robust induction of T helper type 1 and type 2 cytokines. STING agonist ADU-S100 also induced distinct activation with increased secretion of RANTES, MIP-1a, MIP-1b and TNF-α) in all models. Pembrolizumab, magrolimab and their combination showed more modest and selective stimulation in comparison to Immunocult and STING-agonism in each model.

**Figure 3.**
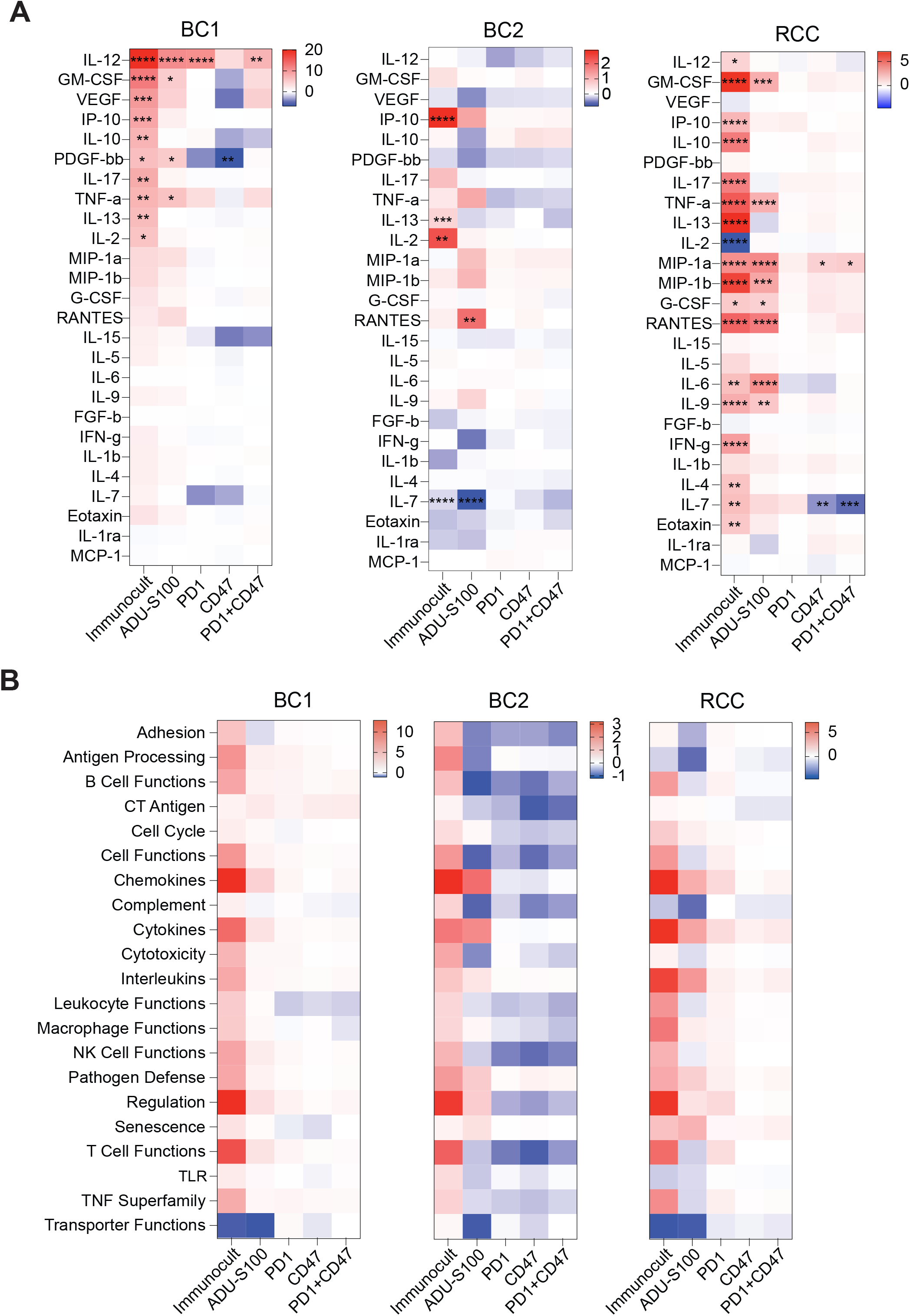
*Ex vivo* cultures reflect biological changes in response to immunomodulation. **A**, BC1 cytokine response **B**, BC2 cytokine response and **C**, RCC cytokine response of each treatment shown as log_2_FoldChange (treated explants to untreated controls). Statistical significance was tested with a two-way ANOVA with Fishers LSD. (**** = p <0.0001; *** = p <0.001; ** = p <0.01; * = p <0.05) **D**, heatmap of average BC1 **E**, BC2 and **F**, RCC patient pathway scores determined by gene expression profiling. The scale reflects the treated pathway score – the untreated pathway score.

All samples in BC1 showed higher response to treatments compared to BC2 indicating these changes may reflect culture conditions rather than tumor material per se. Thus, the immunological responses observed in preclinical *ex vivo* models are more accurately compared within and not across models. Interestingly, IL-2-supplemented RCC showed a significant drop in IL-2 levels following Immunocult treatment **(Fig 3A**), probably reflecting regulatory T cell activation in the model.

### PDECs reflect biological changes in response to immunomodulation

We further confirmed the activation of immune-related pathways using transcriptional profiling with NanoString **(Fig 3B, S6A-D)**. As observed with secreted cytokines, Immunocult led to robust T cell activation in each model with chemokine, cytokine, and interleukin pathway stimulation. This can be interpreted as the maximum T cell activation within the model and can be used as a reference when testing novel immunotherapies **(S6A)**. The STING-agonist ADU-S100 also showed a strong response, although more restricted to expected pathways (eg. chemokines) in innate immune modulation. The RCC samples presented the most notable response to PD-1 blockade in relation to the maximum T cell response observed following Immunocult treatment. The anti-PD-1 response resembled Immunocult response albeit at a lower stimulation magnitude. This was accompanied by an increase in various cell functions (B-, NK- and T-cell, leukocyte, and macrophage functions) as well as stimulation of interleukin, cytokine, and chemokine pathways **(Fig 3B)**. These effects were more modest in the BC models **(Fig 3B)**. However, without clinical response data to anti-PD-1, it is not clear whether the addition of IL-2 in the RCC culture medium primed anti-PD-1 resistant tumor samples to ICB, as seen in *ex vivo* melanoma cultures with low baseline secretion of IL-2^20^, or whether the higher response in comparison to BC is because RCC is generally more sensitive to anti-PD-1 therapy.

We next explored the possibility that tumor properties correlate with *ex vivo* treatment responses. We analyzed the correlation of baseline immune cell abundance with treatment response **(S7A-E)**. For instance, baseline T-cell abundance did not influence the magnitude of the increase in ‘T cell functions’ following treatments aiming at T-cell activation **(S7A)**. Meanwhile, the number of DCs at baseline seemed to closely correlate with macrophage functions following T cell activation, STING agonism, and treatment with anti-PD-1 **(S7A-C)**. Along these lines, other studies have shown that major physiological functions of type I interferons are directed towards DCs^21^, making it plausible that DC numbers correlate with treatment response.

### Core innate and adaptive immune response signatures as potential biomarkers

As BC and RCC *ex vivo* models generally responded similarly to immunomodulation, we pooled the NanoString gene expression analyses from all samples together and generated a core response signature for T cell activation (adaptive, Immunocult) and interferon signaling (innate, ADU-S100). Among the different treatment approaches, Immunocult induced the highest number of differentially expressed genes **(Fig 4A)**, and this was more robustly seen in BC1 and RCC models. The three PDEC models generated a shared upregulated ‘core response’ gene set for Immunocult: *CCL2, CCL7, CCL8, CD274, CFB, CX3CL1, CXCL10, CXCL11, CXCL9, ICAM1, IDO1, IL15RA, IRF1, JAK2, SERPING1, SOCS1, TAP1, TAP2, XCL2* **(Fig 4B)**. The models did not share any downregulated genes following Immunocult treatment **(Fig 4B)**. The pathways associated with the upregulated core response signature were analyzed in the Metascape^22^ validating the responses to the activation of an adaptive immune response and more selectively to type II interferon signaling **(Fig 4C)**. We tested our “adaptive signature” of T cell activation in a dataset of post PD-1-treated melanoma patients, and found that our “T cell signature” was able to stratify patients treated with anti-PD-1^8,9^ for better overall survival on par with existing biomarkers for immunotherapy including IFNg, CD8+ T cell infiltration, and a biomarker developed by Merck **(Fig 4D-E)**.

**Figure 4.**
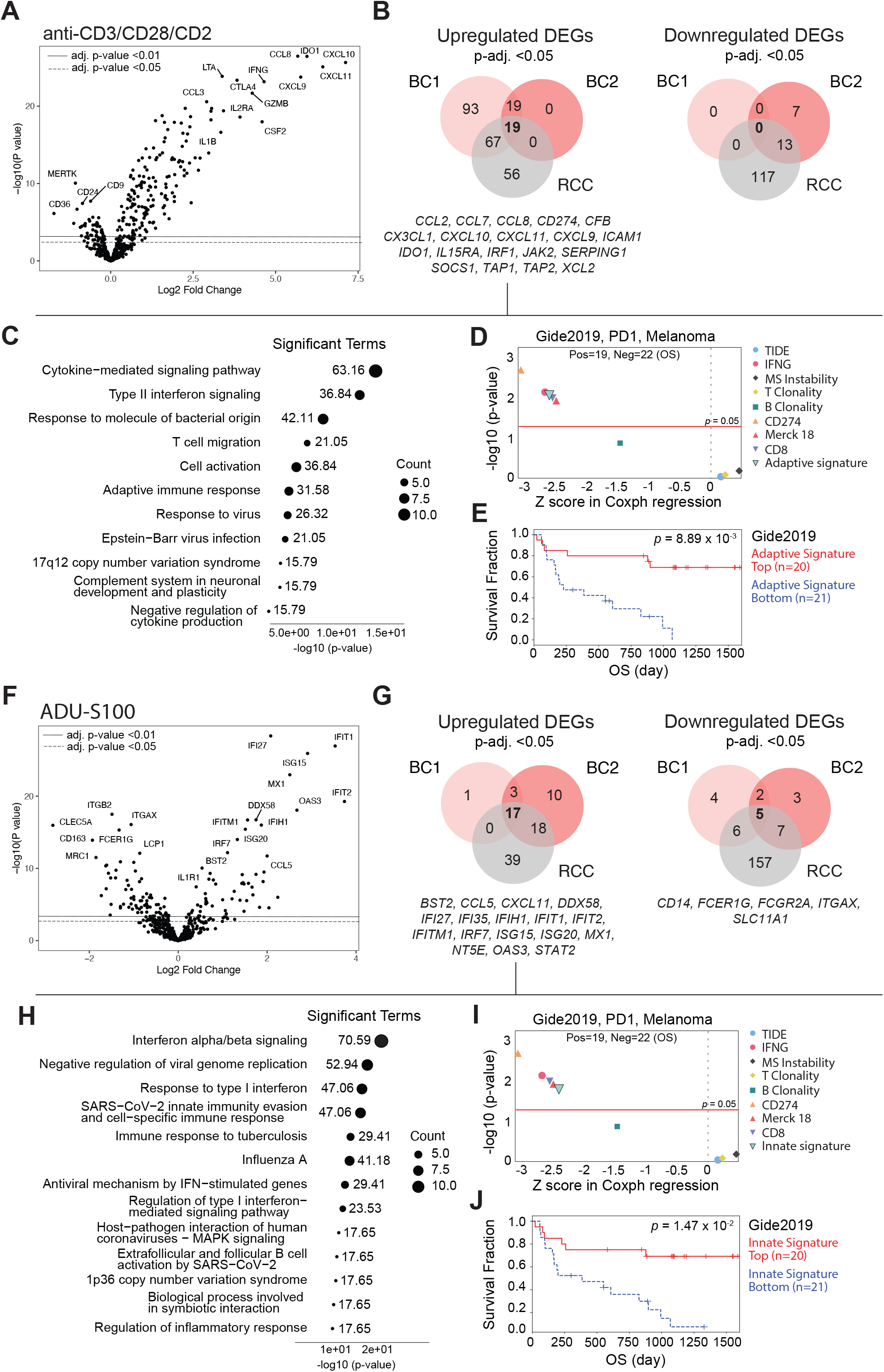
Core innate and adaptive response signatures as potential biomarkers. **A**, Differentially expressed genes of pooled explant samples following Immunocult treatment. **B**, Venn diagrams of significantly (adjusted p-value < 0.5) upregulated and downregulated genes following Immunocult. Shared upregulated core gene signature of 19 genes is bolded, and core genes are listed below the plot. **C**, Metascape analysis of pathways associated with the core signature, circle size refers to the number of core genes (=count) matching the pathway, and the numerical value is the percentage of all of the user-provided genes that are found in the given ontology term. “Log_10_(P)” is the p-value in log base 10. **D**, The core “adaptive signature” in relation to existing biomarkers for immunotherapies in a cohort of anti-PD-1 treated melanoma patients from Gide et al., 2019 **E**, overall survival of anti-PD-1-treated melanoma patients with high (red) vs. low (blue) core “adaptive signature” expression. **F**, Differentially expressed genes of pooled explant samples following ADU-S100 treatment **G**, Venn diagrams of significantly upregulated or downregulated genes following ADU-S100, with core signature genes listed below the plots. **H**, Metascape analysis of pathways associated with the upregulated core signature, circle size refers to the amount of core genes matching the pathway, and the numerical value is the percentage of all of the user-provided genes that are found in the given ontology term. “Log_10_(P)” is the p-value in log base 10. **I**, The core “innate signature” in relation to existing biomarkers for immunotherapies in a set of anti-PD-1 treated melanoma patients **J**, overall survival of anti-PD-1-treated melanoma patients with high expression (red) of the core “innate signature” vs. low expression (blue) of the core “innate signature”. OS, overall survival.

STING-agonism through ADU-S100 induced also robust but more modest gene expression changes than direct T cell activation, as determined by comparing the magnitude of the fold-change response **(Fig 4F)**. The shared upregulated core genes for innate activation were: *BST2, CCL5, CXCL11, DDX58, IFI27, IFI35, IFIH1, IFIT1, IFIT2, IFITM1, IRF7, ISG15, ISG20, MX1, NT5E, OAS3, STAT2* **(Fig 4G)**. Interestingly, BC2 had more unique upregulated genes following ADU-S100 treatment, while BC1 had more upregulated genes following T cell activation **(Fig 4B** and **G)**. Activating the STING pathway has an expected effect of downregulating CD14 which could be seen in all three models **(Fig 4G)**. Upregulated innate core genes were involved in type I interferon response **(Fig 4H)**. This signature was comparable in terms of predicting overall survival as the T cell signature, presumably reflecting the overall positive association of the baseline pro-inflammatory tumor immune microenvironment on ICB clinical response **(Fig 4I-J)**. Individual genes were not significantly upregulated following pembrolizumab, magrolimab, or pembrolizumab + magrolimab treatments **(S8A-C)**. Notably, genes (*CMKLR1, CCL18, IL6R*) within the NF-κB-IL6 pathway were downregulated only with the combination of the two, although not significantly. Targeting this pathway in cancer-associated fibroblasts in BC has been proposed as a way of normalizing the tumor stroma with possible anti-cancer effects^23^.

In summary, our work presents immunotherapy treatment responses in three *ex vivo* models of breast cancer and renal cell carcinoma. We found that T cell activation with Immunocult increased the secretion of cytokines (IP-10, TNF-α) and expression of genes associated with type II interferon signaling, whereas ADU-S100 increased cytokine secretion (RANTES, MIP-1a, MIP-1b and TNF-α) and expression of genes associated with type I signaling in all three models, although in different magnitudes. The shared core adaptive (*CCL2, CCL7, CCL8, CD274, CFB, CX3CL1, CXCL10, CXCL11, CXCL9, ICAM1, IDO1, IL15RA, IRF1, JAK2, SERPING1, SOCS1, TAP1, TAP2, XCL2*) and innate (*BST2, CCL5, CXCL11, DDX58, IFI27, IFI35, IFIH1, IFIT1, IFIT2, IFITM1, IRF7, ISG15, ISG20, MX1, NT5E, OAS3, STAT2)* response signatures and individual patient-specific responses may serve as potential exploratory biomarkers for treatment outcomes.

## Supporting information

SuppFig_and_Captions

## ACKNOWLEDGMENTS

We thank the study participants, the study nurses, the investigators, and the research team who contributed to the study. We also want to thank Mari Parsama and Teija Kanasuo for technical assistance. Nanostring analysis was carried out at the functional genomics unit (FuGU) at the Unviversity of Helsinki.

The present works were mainly funded through the Business Finland Health program award for project Cancer IO. Klefström research group has received funding from the Academy of Finland, Business Finland, the Finnish Cancer Organizations, Sigrid Juselius foundation, Jane and Aatos Erkko foundation, the Research Council of Finland, and RESCUER project, which has received funding from the European Union’s Horizon 2020 Framework Programme (no. 847912). This work was also supported by the U.S. Department of Defense for Health Affairs through the Breast Cancer Research Program (award no. W81XWH2110773). Opinions, interpretations, conclusions, and recommendations are those of the author and are not necessarily endorsed by the Department of Defense. In addition, funds were received from Sihtasutus Archimedes, Ida Montinin Säätiö, Finnish Cancer Institute, Syöpäjärjestöt, and the iCAN Digital Precision Cancer Medicine Flagship. Mustjoki research group has received funding from Cancer Foundation Finland, Academy of Finland, Sigrid Juselius Foundation, Gyllenberg Foundation, Jane and Aatos Erkko Foundation, State funding for the University-level Health Research in Finland, and HiLIFE fellow funds. Hollmén research group has received funding from Academy of Finland, Cancer Foundations, and the Sigrid Jusélius Foundation.

## AUTHOR CONTRIBUTIONS

R.T., A.T., J.P., M.H., S.M., and J.K., designed the study; R.T, K.P., J.H.R., R.L, and A.N.K optimized *ex vivo* cultures and treated explants, and collected data. R.T., K.P., J.H.R., and D.N analyzed and visualized the data. M.M., P.E.K., L.N., T.M., J.M., P.J., K.L., R.J., S.T., T.M., P.B., and I.K., recruited patients, and collected clinical data, R.T., K.P., M.H., S.M, and J.K interpreted the data, R.T, K.P., M.H., S.M., J.K., wrote the first draft of the manuscript. All authors reviewed the manuscript and approved the final version.

## REFERENCES

1. Najafi, M. et al. Tumor microenvironment: Interactions and therapy. Journal of Cellular Physiology vol. 234 Preprint at 10.1002/jcp.27425 (2019).

2. Voabil, P. et al. An ex vivo tumor fragment plaRorm to dissect response to PD-1 blockade in cancer. Nat Med 27, (2021).

3. Yuki, K., Cheng, N., Nakano, M. & Kuo, C. J. Organoid Models of Tumor Immunology. Trends in Immunology vol. 41 Preprint at 10.1016/j.it.2020.06.010 (2020).

4. Jenkins, R. W. et al. Ex Vivo Profiling of PD-1 Blockade Using Organotypic Tumor Spheroids. (2018) doi:10.1158/2159-8290.CD-17-0833.

5. Turpin, R. et al. Respiratory complex I regulates dendritic cell maturation in explant model of human tumor immune microenvironment. J Immunother Cancer 12, (2024).

6. Munne, P. M. et al. Compressive stress-mediated p38 activation required for ERα + phenotype in breast cancer. Nat Commun 12, (2021).

7. Benjamini, Y. & Hochberg, Y. Controlling the False Discovery Rate: A Practical and Powerful Approach to Multiple Testing. Journal of the Royal Sta=s=cal Society: Series B (Methodological) 57, (1995).

8. Jiang, P. et al. Signatures of T cell dysfunction and exclusion predict cancer immunotherapy response. Nat Med 24, (2018).

9. Fu, J. et al. Large-scale public data reuse to model immunotherapy response and resistance. Genome Med 12, (2020).

10. Thorsson, V. et al. The Immune Landscape of Cancer. Immunity 48, (2018).

11. Shankar, S. P. Dendritic cells and the extracellular matrix: A challenge for maintaining tolerance/homeostasis. World J Immunol 5, (2015).

12. Hollmén, M. et al. G-CSF regulates macrophage phenotype and associates with poor overall survival in human triple-negative breast cancer. Oncoimmunology 5, (2016).

13. Rugo, H. S. et al. Abstract GS3-01: Additional efficacy endpoints from the phase 3 KEYNOTE-355 study of pembrolizumab plus chemotherapy vs placebo plus chemotherapy as first-line therapy for locally recurrent inoperable or metastatic triple-negative breast cancer. Cancer Res 81, (2021).

14. Cortés, J. et al. LBA16 KEYNOTE-355: Final results from a randomized, double-blind phase III study of first-line pembrolizumab + chemotherapy vs placebo + chemotherapy for metastatic TNBC. Annals of Oncology 32, (2021).

15. Rini, B. I. et al. Pembrolizumab plus Axitinib versus Sunitinib for Advanced Renal-Cell Carcinoma. New England Journal of Medicine 380, (2019).

16. Choueiri, T. K. et al. Adjuvant Pembrolizumab amer Nephrectomy in Renal-Cell Carcinoma. New England Journal of Medicine 385, (2021).

17. Motzer, R. J. et al. Nivolumab versus Everolimus in Advanced Renal-Cell Carcinoma. New England Journal of Medicine 373, (2015).

18. Xu, J. X. et al. FDA Approval Summary: Nivolumab in Advanced Renal Cell Carcinoma Amer Anti-Angiogenic Therapy and Exploratory Predictive Biomarker Analysis. Oncologist 22, (2017).

19. Park, H. R. et al. Blockade of CD47 enhances the antitumor effect of macrophages in renal cell carcinoma through trogocytosis. Sci Rep 12, (2022).

20. Kaptein, P. et al. Addition of interleukin-2 overcomes resistance to neoadjuvant CTLA4 and PD1 blockade in ex vivo patient tumors. Sci Transl Med 14, (2022).

21. Diamond, M. S. et al. Type I interferon is selectively required by dendritic cells for immune rejection of tumors. Journal of Experimental Medicine 208, (2011).

22. Zhou, Y. et al. Metascape provides a biologist-oriented resource for the analysis of systems-level datasets. Nat Commun 10, (2019).

23. Hendrayani, S. F., Al-Harbi, B., Al-Ansari, M. M., Silva, G. & Aboussekhra, A. The inflammatory/cancer-related IL-6/STAT3/NF-κB positive feedback loop includes AUF1 and maintains the active state of breast myofibroblasts. Oncotarget 7, (2016).

