## Supplementary material for "Patient-derived tumor explant models of tumor immune microenvironment reveal distinct and reproducible immunotherapy responses": SuppFig_and_Captions

Turpin et al., Supplementary Figure 1.

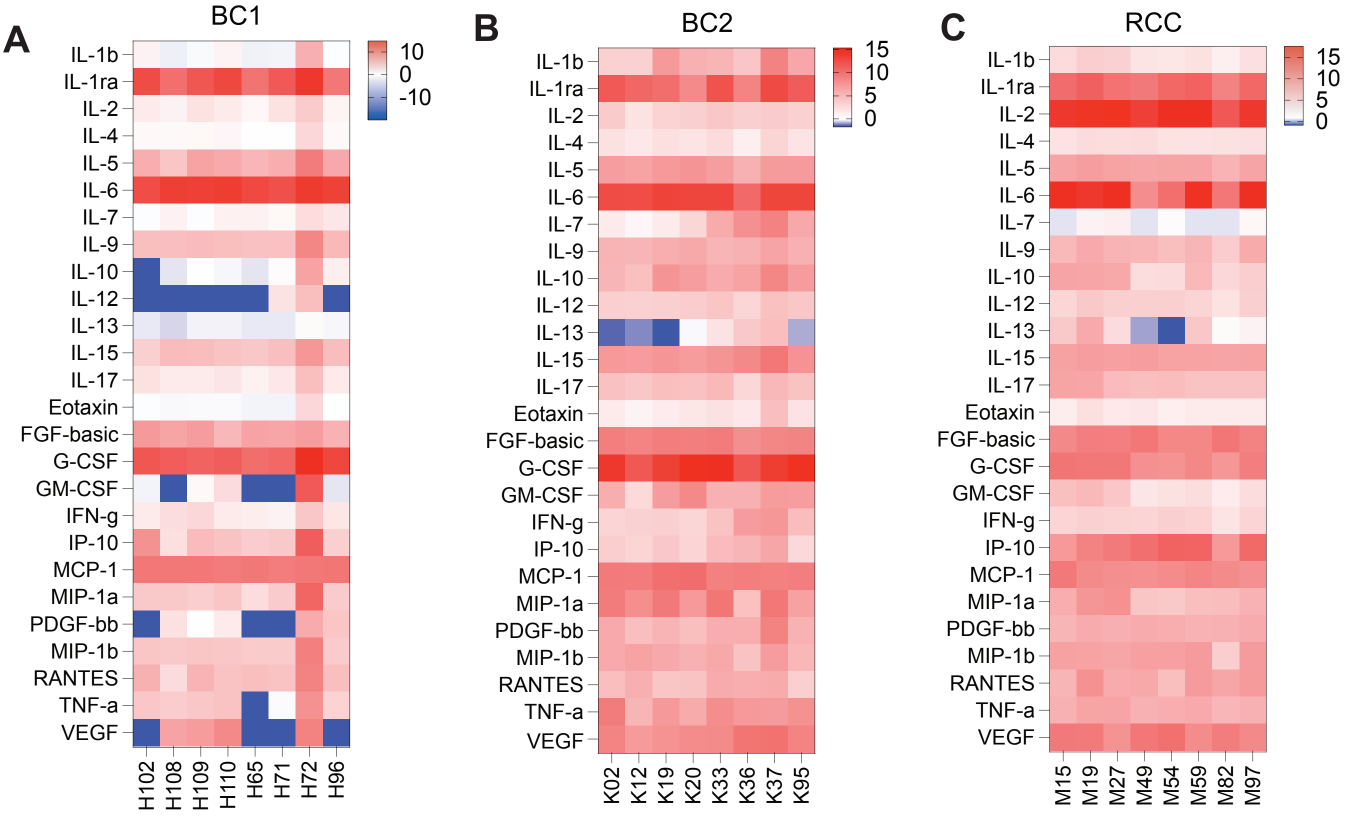


D

Breast cancer

| | |
|---|---|
| H65 | Luminal B |
| H72 | TNBC |
| H96 | Luminal B |
| H108 | Luminal A |
| H109 | Luminal B |
| H110 | Luminal B |
| K95 | Luminal B |
| K02 | Luminal B |
| K12 | Luminal B |
| K20 | DCIS |
| K33 | Luminal B |
| K37 | Luminal B |

RCC

| | |
|---|---|
| M59 | Clear cell |
| M15 | Chromophobic |
| M97 | Clear cell |
| M49 | Clear cell |
| M54 | Clear cell |
| M82 | Clear cell + papillary |

**Supplementary Figure 1. Individual patient cytokine profiles and clinical diagnosis A,** Individual patient baseline cytokine secretion after 48hrs for BC1 **B,** BC2, **C,** RCC. Scale shows $\log_2$ of the raw pg/ml (+1 as a small constant value).
**D,** clinical subtypes of BC1, BC2 and RCC patients.

Turpin et al., Supplementary Figure 2.

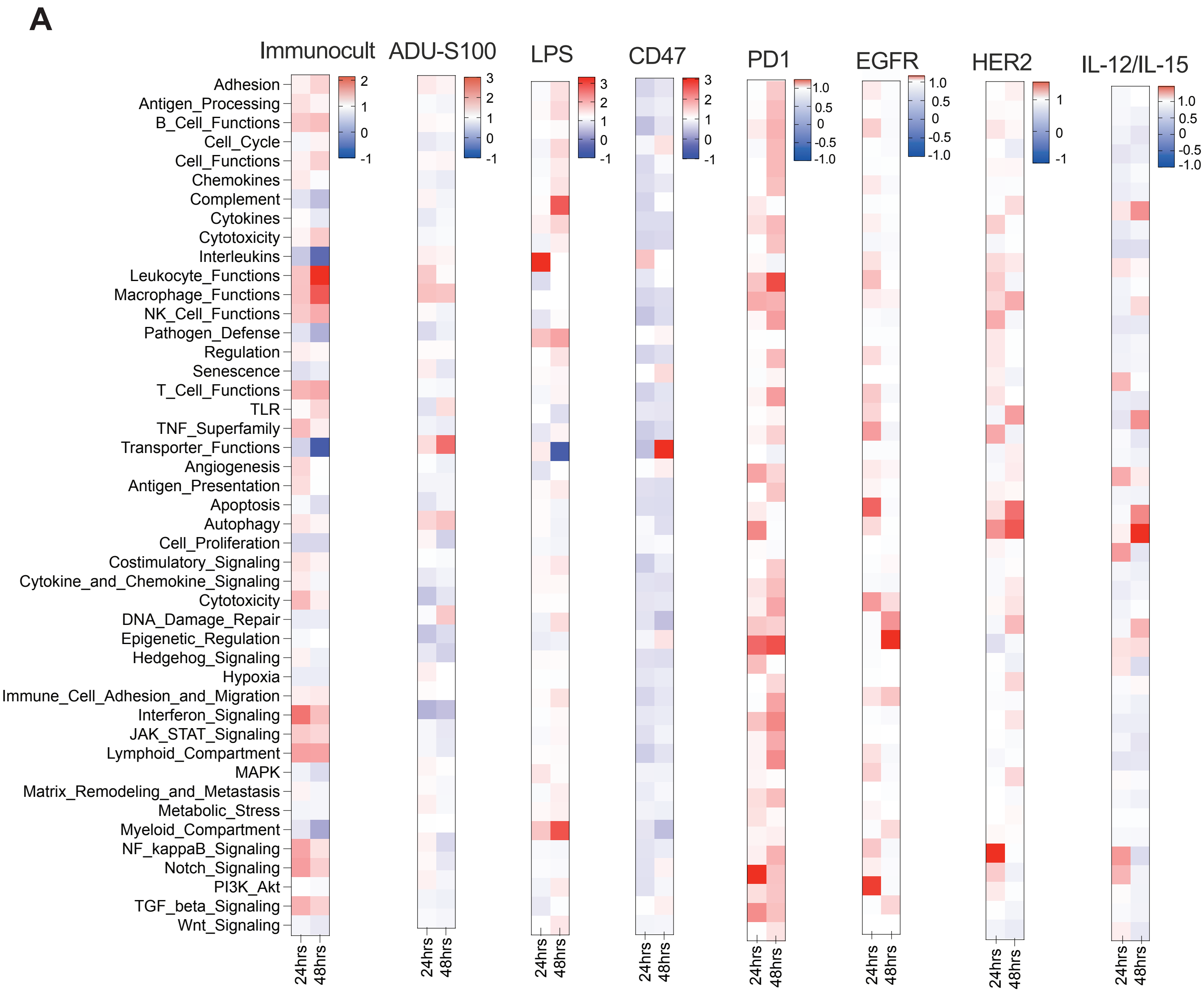

**Supplementary Figure 2. Pilot pathway analysis based on gene expression profiling A,** RCC patient explant 24hrs and 48hrs following Immunocult**.** BC2 patient explant 24hrs and 48hrs following ADU-S100**.** BC1 patient explant 24hrs and 48hrs following LPS**.** BC1 patient explant 24hrs and 48hrs following anti-CD47**.** RCC patient explant 24hrs and 48hrs following anti-PD-1**.** RCC patient explant 24hrs and 48hrs following cetuximab (EGFR**).** BC2 patient explant 24hrs and 48hrs following trastuzumab (HER2)**.** BC2 patient explant 24hrs and 48hrs following IL-2/IL-15. $Log_2$FoldChanges compared to untreated sample of the corresponding timepoint (24hrs treated/control, 48hrs treated/control).

Turpin et al., Supplementary Figure 3.

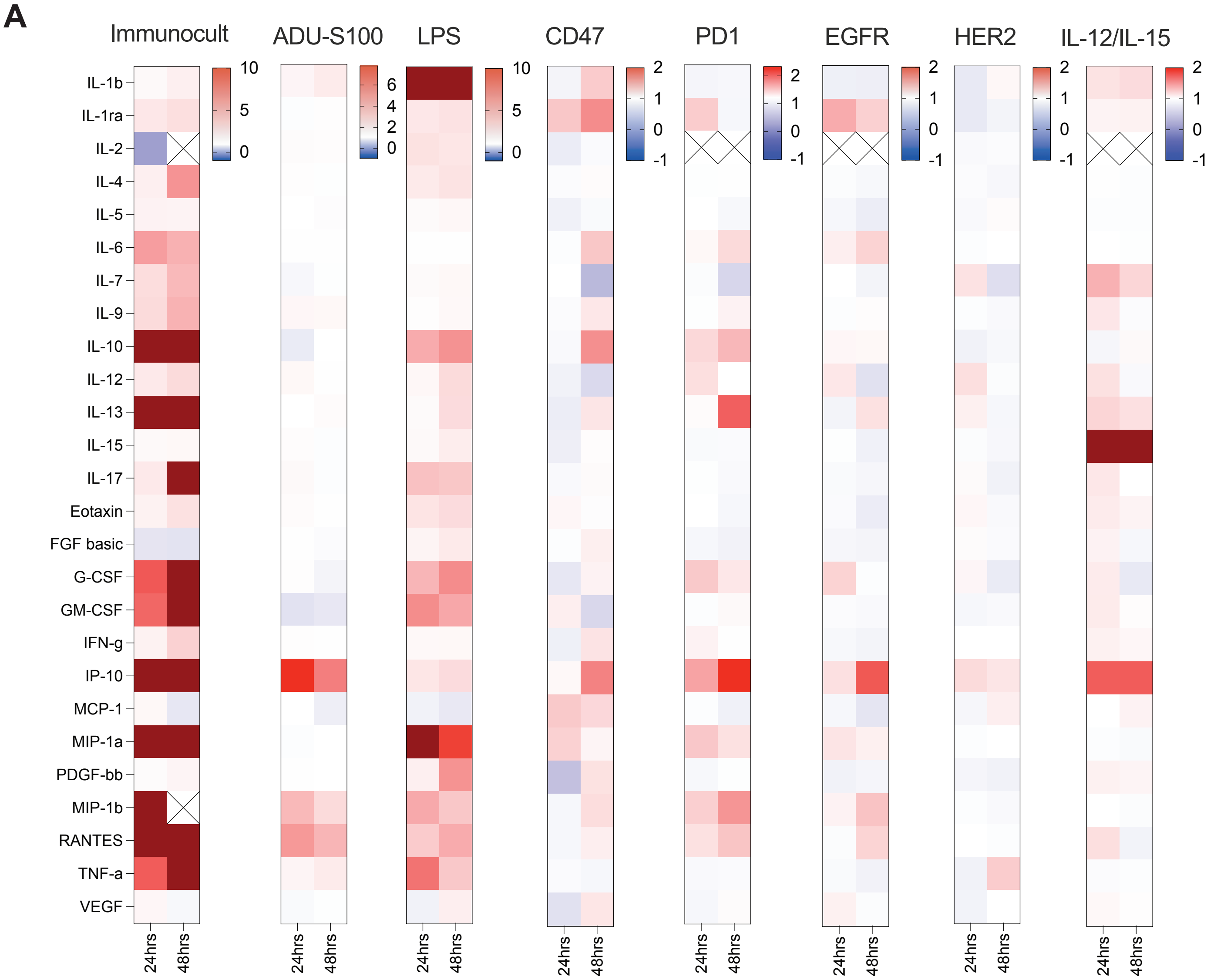

**Supplementary Figure 3. Pilot cytokine profiling A,** RCC patient explant 24hrs and 48hrs following Immunocult**.** BC2 patient explant 24hrs and 48hrs following ADU-S100**.** BC1 patient explant 24hrs and 48hrs following LPS**.** BC1 patient explant 24hrs and 48hrs following anti-CD47**.** RCC patient explant 24hrs and 48hrs following anti-PD-1**.** RCC patient explant 24hrs and 48hrs following cetuximab (EGFR)**.** BC2 patient explant 24hrs and 48hrs following trastuzumab (HER2)**.** BC2 patient explant 24hrs and 48hrs following IL-2/IL-15. $Log_2$FoldChanges compared to control sample of the corresponding timepoint (24hrs treated/control, 48hrs treated/control). Blank and excluded values marked with an X, dark red squares above defined range.

Turpin et al., Supplementary Figure 4.

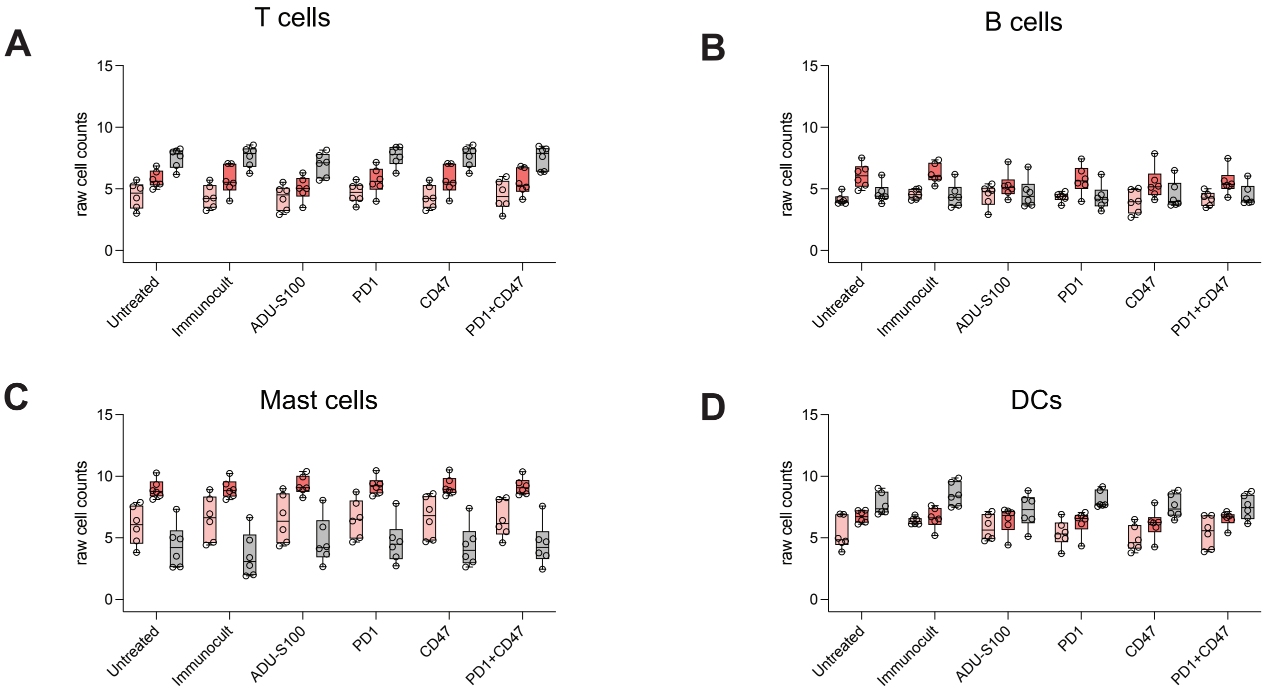

**Supplementary Figure 4. Cell types not significantly affected by immunomodulation A-D,** T cell, B cell, mast cell and DC numbers in culture determined by gene expression profiling following treatment. Data shown as box and whisker plot min to max with a line at the median. Statistics were calculated by 2-way ANOVA with Fishers exact test.

Turpin et al., Supplementary Figure 5.

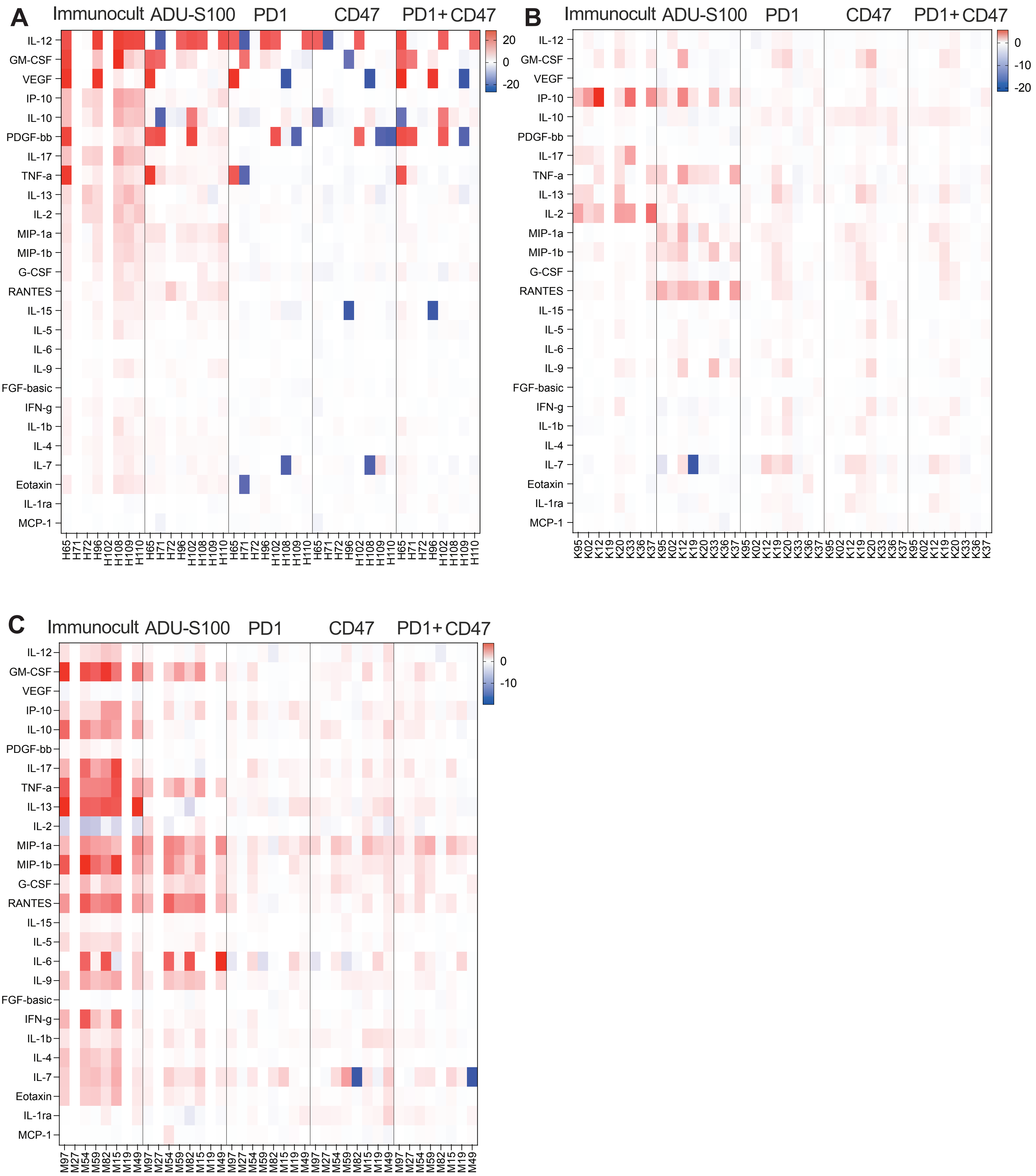

**Supplementary Figure 5. Individual patient cytokine responses to immunomodulation A,** individual patient BC1 cytokine profiles in response to treatments **B,** individual patient BC2 cytokine profiles in response to treatments **C,** individual patient RCC cytokine profiles in response to indicated treatments. Scale shows $log_2$FoldChanges (treated/untreated).

Turpin et al., Supplementary Figure 6.

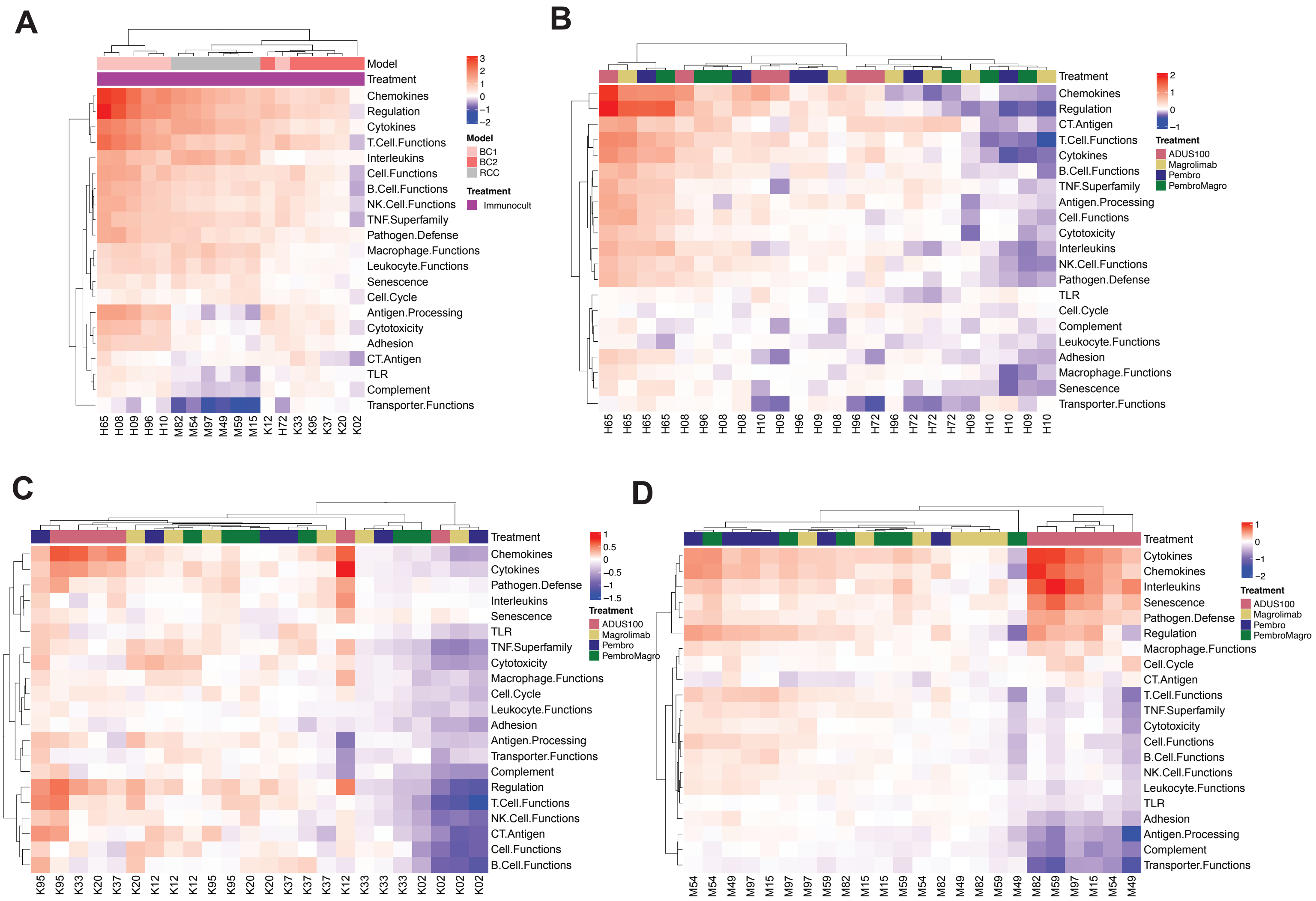

**Supplementary Figure 6. Individual patient pathway analysis in response to immunomodulation. A,** unsupervised clustering of individual patient pathway score changes in response to Immunocult treatment in all models. **B-D,** unsupervised clustering of ADU-S100 treatment, anti-PD-1 treatment, anti-CD47 treatment, and anti-PD-1 + anti-CD47 treatment pathway scores in **B,** BC1, **C,** BC2, and **D,** RCC models. Pathway scores displayed as $\log_2$(treated score+7) - $\log_2$(untreated score+7).

Turpin et al., Supplementary Figure 7.

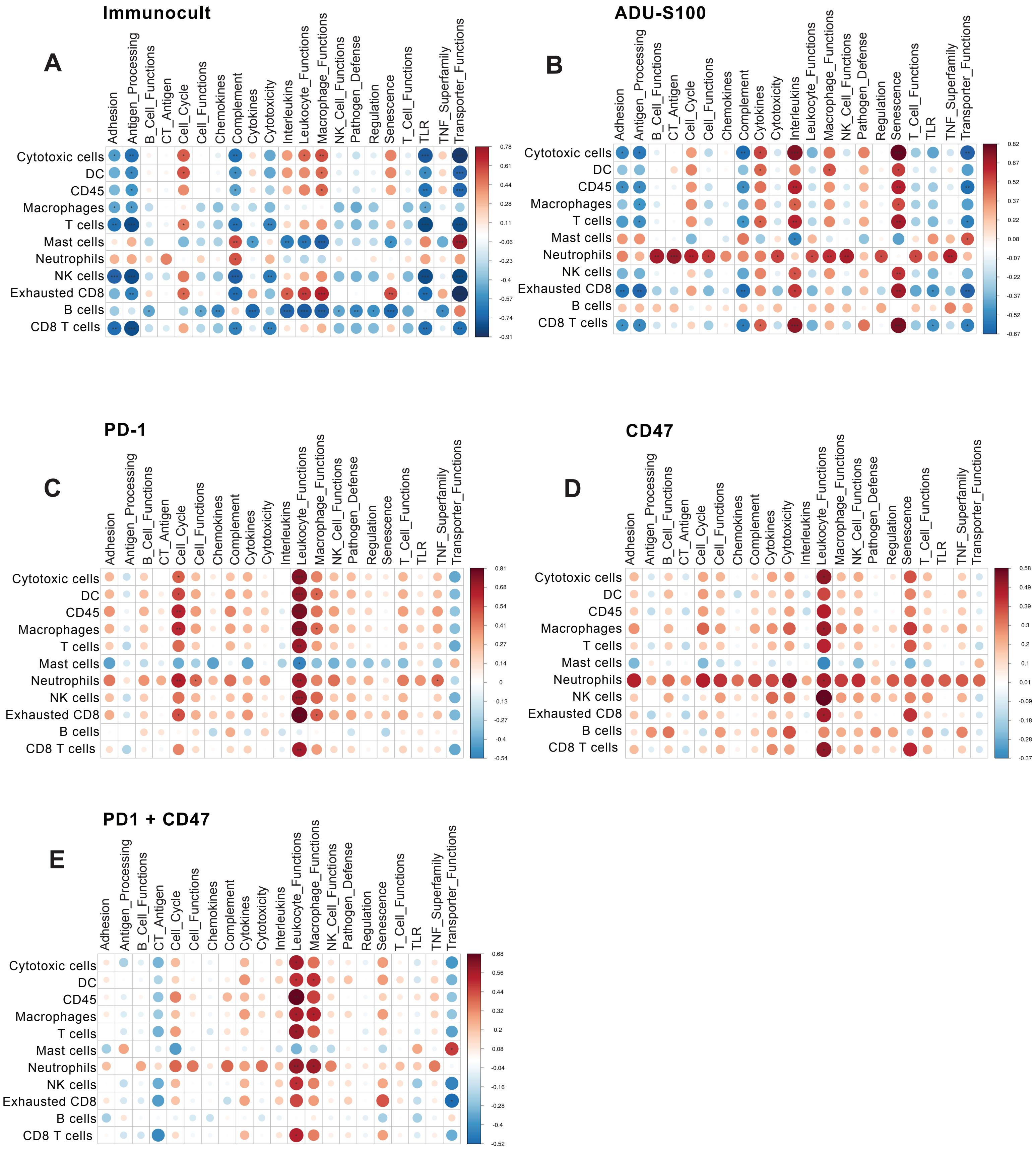

**Supplementary Figure 7. Correlation plots comparing baseline immune cell amounts to treatment response. A - E,** Pearson correlation of the average baseline cell type score compared to the magnitude of the treatment response to **A,** Immunocult **B,** ADU-S100 **C,** anti-PD-1 **D,** anti-CD47 **E,** or anti-PD-1 + anti-CD47. Circle size and color scales refer to Pearson correlation coefficient values.

Turpin et al., Supplementary Figure 8.

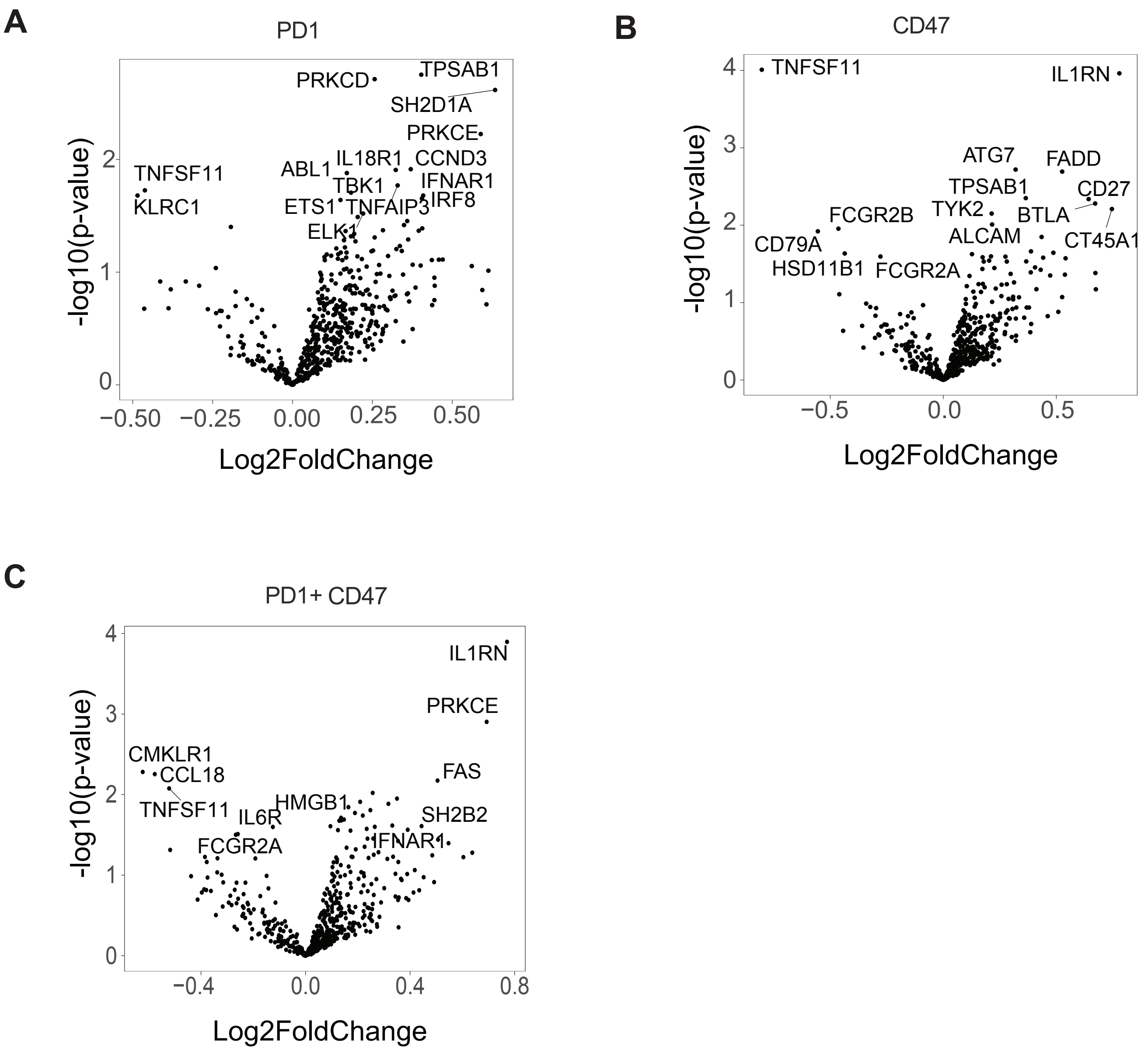

**Supplementary Figure 8. Volcano plots displaying differentially expressed genes in pooled patient explant samples. A - C,** Volcano plot of differentially expressed genes following anti-PD-1 (A), anti-CD47 (B) or anti-PD-1 + anti-CD47 (C) treatment. Adjusted p-value is not shown as none of the data points have reached statistical significance.
